# An updated sulfate transporter phylogeny uncovers a perennial-specific subgroup associated with lignification

**DOI:** 10.1101/2025.04.23.650276

**Authors:** Samantha M. Surber, Chen Hsieh, Lan Na, Scott A. Harding, Chung-Jui Tsai

**Affiliations:** Department of Plant Biology, University of Georgia, Athens, GA 30602, USA; Center for Bioenergy Innovation, Oak Ridge National Laboratory, Oak Ridge, TN 37831, USA; Institute of Bioinformatics, University of Georgia, Athens, GA 30602, USA; Complex Carbohydrate Research Center, University of Georgia, Athens, GA 30602, USA; Warnell School of Forestry, University of Georgia, Athens, GA 30602, USA; Department of Genetics, University of Georgia, Athens, GA 30602, USA

**Keywords:** SULTR, Populus, sulfur assimilation, lignin biosynthesis, xylem

## Abstract

Sulfate-proton co-transporters (SULTRs) mediate sulfate uptake, transport, storage, and assimilation in plants. The SULTR family has historically been classified into four groups (SULTR1-SULTR4), with well-characterized roles for SULTR groups 1, 2, and 4. However, the functions of the large and diverse SULTR3 group remain poorly understood. Here, we present an updated phylogenetic analysis of SULTRs across angiosperms, including multiple early-divergent lineages. Our results suggest that the enigmatic SULTR3 group comprises four distinct subfamilies that predate the emergence of angiosperms, providing a basis for reclassifying the SULTR family into seven subfamilies. This expanded classification is supported by subfamily-specific gene structures and amino acid substitutions in the substrate-binding pocket. Structural modeling identified three serine residues uniquely lining the substrate-binding pocket of SULTR3.4, enabling three hydrogen bonds with the phosphate ion. The data support the proposed neofunctionalization of this subfamily for phosphate allocation within vascular tissues. Transcriptome analysis of *Populus tremula* × *alba* revealed divergent tissue expression preferences among *SULTR* subfamilies and between genome duplicates. We observed partitioned expression in vascular tissues among the four *SULTR3* subfamilies, with *PtaSULTR3.4a* and *PtaSULTR3.2a* preferentially expressed in primary and secondary xylem, respectively. Gene coexpression analysis revealed coordinated expression of *PtaSULTR3.4a* with genes involved in phosphate starvation responses and nutrient transport, consistent with a potential role in phosphate homeostasis. In contrast, *PtaSULTR3.2a* was strongly coexpressed with lignification and one-carbon metabolism genes and their upstream transcription regulators. *PtaSULTR3.2a* belongs to a eudicot-specific branch of the SULTR3.1 subfamily found only in perennial species, suggesting a specialized role in lignifying tissues. Together, our findings provide a refined phylogenetic framework for the SULTR family and suggest that the expanded SULTR3 subfamilies have undergone neofunctionalization during the evolution of vascular and perennial plants.

## INTRODUCTION

Sulfur is an essential plant macronutrient taken up from the soil as inorganic sulfate (Kopriva et al. 2019). Uptake and plantwide distribution are mediated by a family of <u>sul</u>fate-proton co-<u>tr</u>ansporters (SULTRs) historically classified into four groups, referred to as SULTR1 to SULTR4 (Takahashi et al. 2000). This Arabidopsis (*Arabidopsis thaliana*)-based nomenclature has been followed in many subsequent phylogenetic analyses (Takahashi et al. 2012; Gallardo et al. 2014; Ding et al. 2016; Xun et al. 2021; Zhao et al. 2022; Chen et al. 2024). Two recent studies classified the SULTR family into five groups but with inconsistent nomenclature and memberships (Heidari et al. 2023; Puresmaeli et al. 2023). Meanwhile, molecular characterization continues to shed light on all SULTR groups as evidenced by their varied subcellular localization, transport kinetics, tissue expression, and stress responsiveness (Takahashi 2019).

Among the different groups, SULTR4 transporters are tonoplast-localized and facilitate sulfate efflux from the vacuole (Kataoka et al. 2004b). SULTR1 and SULTR2 members are plasma membrane-localized with SULTR1 being high-affinity transporters involved in sulfate uptake by the roots (Takahashi et al. 2000; Yoshimoto et al. 2002), and SULTR2 being low-affinity transporters expressed primarily in the root cylinder for root-to-shoot sulfate transport (Takahashi et al. 1997; Takahashi et al. 2000). SULTR3 is the largest and most diverse group, with five genes in Arabidopsis—*AtSULTR3.1, AtSULTR3.2, AtSULTR3.3, AtSULTR3.4*, and *AtSULTR3.5* (Zuber et al. 2010; Takahashi et al. 2012). The subcellular distribution and function of SULTR3 members are less precisely known but recent reports suggest localization to the plasma membrane (Kataoka et al. 2004a; Yamaji et al. 2017; Ding et al. 2020), endoplasmic reticulum (Zhao et al. 2016), and/or chloroplast (Cao et al. 2013; Chen et al. 2019).

SULTR3 members are enigmatic with respect to their sulfate transport capabilities. AtSULTR3.1, AtSULTR3.2, and AtSULTR3.3 did not show sulfate uptake activity in yeast (Takahashi et al. 2000). To date, successful restoration of yeast mutant growth on sulfate-containing media by SULTR3 has only been reported for LjSST1 (SULTR3.5) of lotus (*Lotus japonicus*) and MhSULTR3.1a of apple (*Malus hupehensis*), although the uptake kinetics were not determined (Krusell et al. 2005; Xun et al. 2021). Cao et al. (2013) used individual *sultr3* mutants to show that all Arabidopsis SULTR3 members, except AtSULTR3.5, contribute substantially to sulfate uptake by isolated chloroplasts. Although AtSULTR3.5 alone does not support sulfate uptake by isolated Arabidopsis chloroplasts or in yeast, its co-expression with *AtSULTR2.1* improved growth of yeast mutants over those expressing *AtSULTR2.1* alone (Kataoka et al. 2004a). This cooperation of AtSULTR3.5 and AtSULTR2.1 in the vasculature contributes to root-to-shoot sulfate transport in Arabidopsis (Kataoka et al. 2004a). In rice (*Oryza sativa*), two independent *low phytic acid* (*lpa*) mutant alleles were both mapped to *OsSULTR3.3*, even though no *in vitro* transport activity of OsSULTR3.3 for sulfate, phosphate, inositol, or inositol triphosphate could be demonstrated in yeasts or *Xenopus* oocytes (Zhao et al. 2016). Similarly, SULTR3.4 was independently characterized as a SULTR-like phosphorus distribution transporter (SPDT) involved in phosphate allocation from xylem to developing tissues in rice, barley (*Hordeum vulgare*), and Arabidopsis (Yamaji et al. 2017; Ding et al. 2020; Gu et al. 2022). OsSULTR3.4, HvSULTR3.4, and AtSULTR3.4 all function as an influx transporter for phosphate, but not sulfate, in oocytes, and their knockout mutants showed impaired phosphorus distribution to developing organs (Yamaji et al. 2017; Ding et al. 2020; Gu et al. 2022). These studies highlight some of the ongoing challenges in elucidating the evolution and function of the diverse SULTR3 group.

*SULTR* gene expression is sensitive to abiotic stressors, including drought and salt, in Arabidopsis (Cao et al. 2014), barrelclover (*Medicago truncatula*) (Gallardo et al. 2014), poplar (*Populus trichocarpa*) (Malcheska et al. 2017), maize (*Zea mays*) (Huang et al. 2018), wheat (*Triticum turgidum* L. ssp. *durum*)(Puresmaeli et al. 2023), and soybean (*Glycine max*) (Zhou et al. 2024). The responsiveness of *SULTR* genes to drought and other abiotic stresses varies between studies in part because expression is modulated not only by plant sulfur status (Vidmar et al. 1999; Takahashi et al. 2000; Rouached et al. 2008; Maruyama-Nakashita et al. 2015), but also by stress hormones like ABA (abscisic acid) and MeJA (methyl jasmonic acid) (Cao et al. 2014; Xun et al. 2021). Notably, four of five *AtSULTR3* genes were identified among the 124 transporter transcripts enriched in guard cells (Bauer et al. 2013). Moreover, quintuple *sultr3* mutants showed impaired sulfate uptake for stress-induced ABA synthesis in Arabidopsis (Chen et al. 2019). This supports sulfate as a xylem-borne signal that precedes localized ABA biosynthesis and stomatal closure in response to drought (Ernst et al. 2010; Malcheska et al. 2017; Batool et al. 2018).

Studies on gene expression and the seasonal dynamics of wood and xylem sap sulfate abundance in *Populus tremula* × *alba* have provided additional insights into the function of SULTR3 in stems. The transcripts of *PtaSULTR3.1b, PtaSULTR3.2a*, and *PtaSULTR3.3a* peaked during active growth in the summer (Dürr et al. 2010; Ko et al. 2011; Malcheska et al. 2013). The hydraulic transport network for long-distance and lateral movements of solutes and water are more complex in large stature trees (Pfautsch et al. 2015). In addition, the formation of large woody sinks depends heavily on sulfur assimilation to supply one-carbon (C1) units for methyl-intensive lignification (Rajinikanth et al. 2007).

Here, we report an updated SULTR phylogeny using 20 angiosperm species, including several woody perennials where SULTR function has been little explored. The inclusion of basal lineages revealed a much more ancient origin of the expanded SULTR3 group than previously recognized. Multiple lines of evidence, including gene structure, protein structural modeling with differential substrate docking, and gene expression, support the functional divergence of SULTR3. We also discuss the potential association of certain SULTR3 members with lignification in *Populus*.

## METHODS

### Sequence collection and annotation

SULTR protein sequences from 22 species were obtained from Phytozome v13 (https://phytozome-next.jgi.doe.gov/)(Goodstein et al. 2012), and the list, including gene models, genome version, and species, is provided in Supplemental Table S1. Sequences were aligned using the multiple sequence alignment tool, T-Coffee (Notredame et al. 2000) for inspection. Sequences with large gaps, misalignment, and/or gene structure misannotation were manually curated, sometimes in consultation with independently annotated genome versions at NCBI. The full sequence dataset has been deposited with the U.S. Department of Energy’s Office of Scientific and Technical Information (OSTI.gov). Exon-intron annotation was retrieved from Phytozome for spikemoss (*Selaginella moellendorffii*), moss (*Physcomitrium patens*), pineapple (*Ananas comosus*), blue columbine (*Aquilegia coerulea*), Arabidopsis (*Arabidopsis thaliana*), poplar (*Populus trichocarpa*), rice (*Oryza sativa*), and sorghum (*Sorghum bicolor*) and the structure was illustrated using Gene Structure Draw Tool (https://www.bioinformatics.uni-muenster.de/tools/strdraw/).

### Phylogenetic reconstruction

Preliminary alignment of all 262 protein sequences performed with T-Coffee on EMBL-EBI (Madeira et al. 2024) was used in a model test with ModelFinder (Kalyaanamoorthy et al. 2017) implemented in IQ-TREE v1.6.12 (Nguyen et al. 2015). This revealed a general JTT model for amino acid substitution rate (Jones et al. 1992), gamma rate of 4 (G4), allowing for a proportion of invariable sites (I), and empirical frequencies (F) [JTT+G4+I+F]. The model JTT+G4+I+F was used for maximum likelihood inference for phylogenetic tree construction with IQ-TREE v1.6.12 and 1000 ultrafast bootstrap cycles. Trees were visualized on TreeViewer (https://treeviewer.org/) (Bianchini and Sánchez-Baracaldo 2024) and edited in MEGA-X (Kumar et al. 2018) for presentation. The full tree (Figure 1) is unrooted, while the SULTR3-specific tree (Figure 2) is rooted on the basal spikemoss SULTR3 sequences.

**Figure 1.**
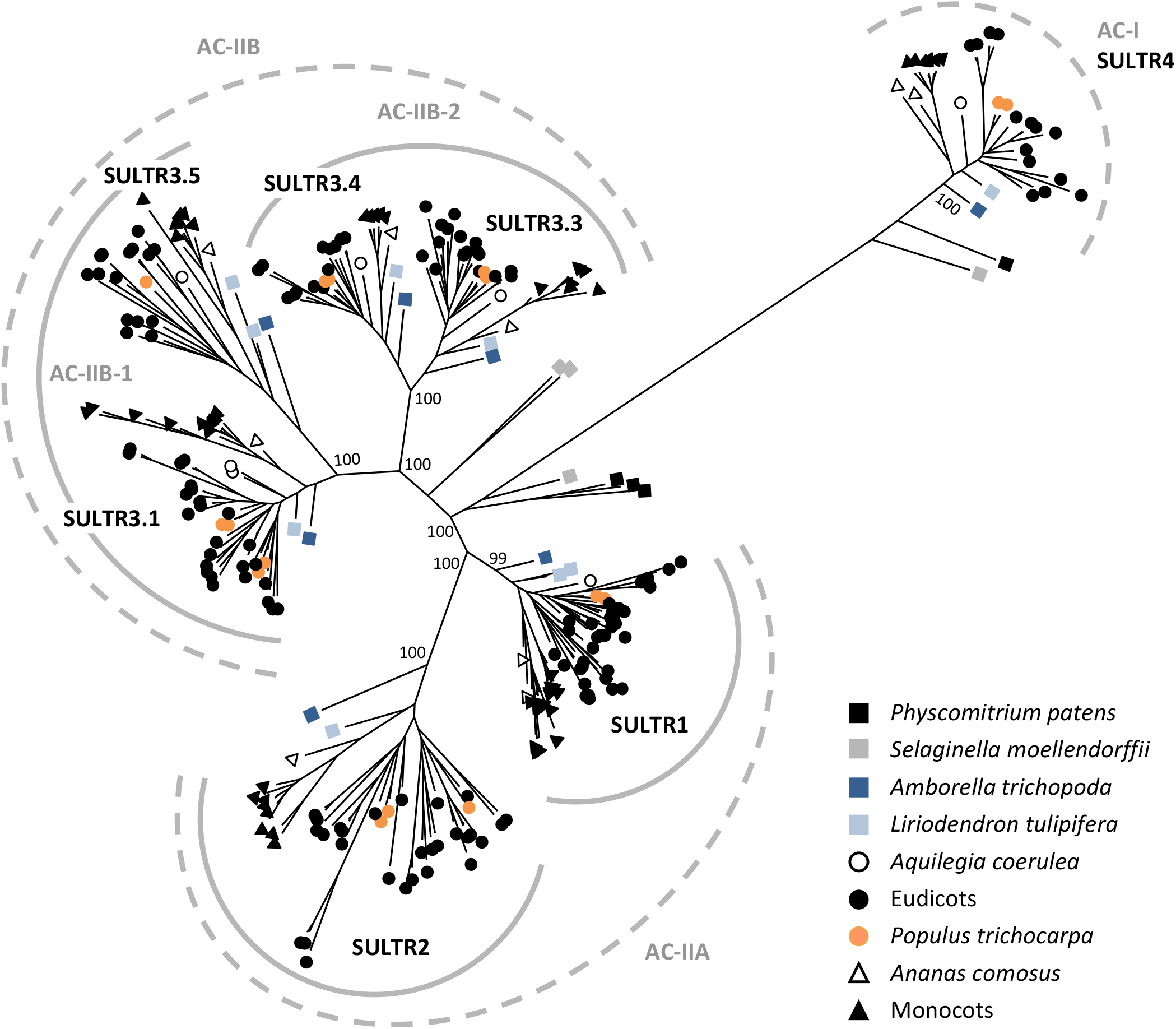
Maximum likelihood phylogenetic inference of 262 SULTR sequences from 22 plant genomes. Angiosperm SULTRs form two ancient clades, AC-I and AC-II. The AC-I clade represents the SULTR4 subfamily. The AC-II clade consists of two subclades, with AC-IIA represented by the SULTR1 and SULTR2 subfamilies, and AC-IIB by SULTR3. AC-IIB is present exclusively in vascular plants with four well-supported groups organized into two branches in all angiosperm taxa. The AC-IIB-1 branch contains SULTR3.1 and SULTR3.5 subfamilies, whereas AC-IIB-2 includes SULTR3.3 and SULTR3.4 subfamilies. Bootstrap values for major nodes are shown.

**Figure 2.**
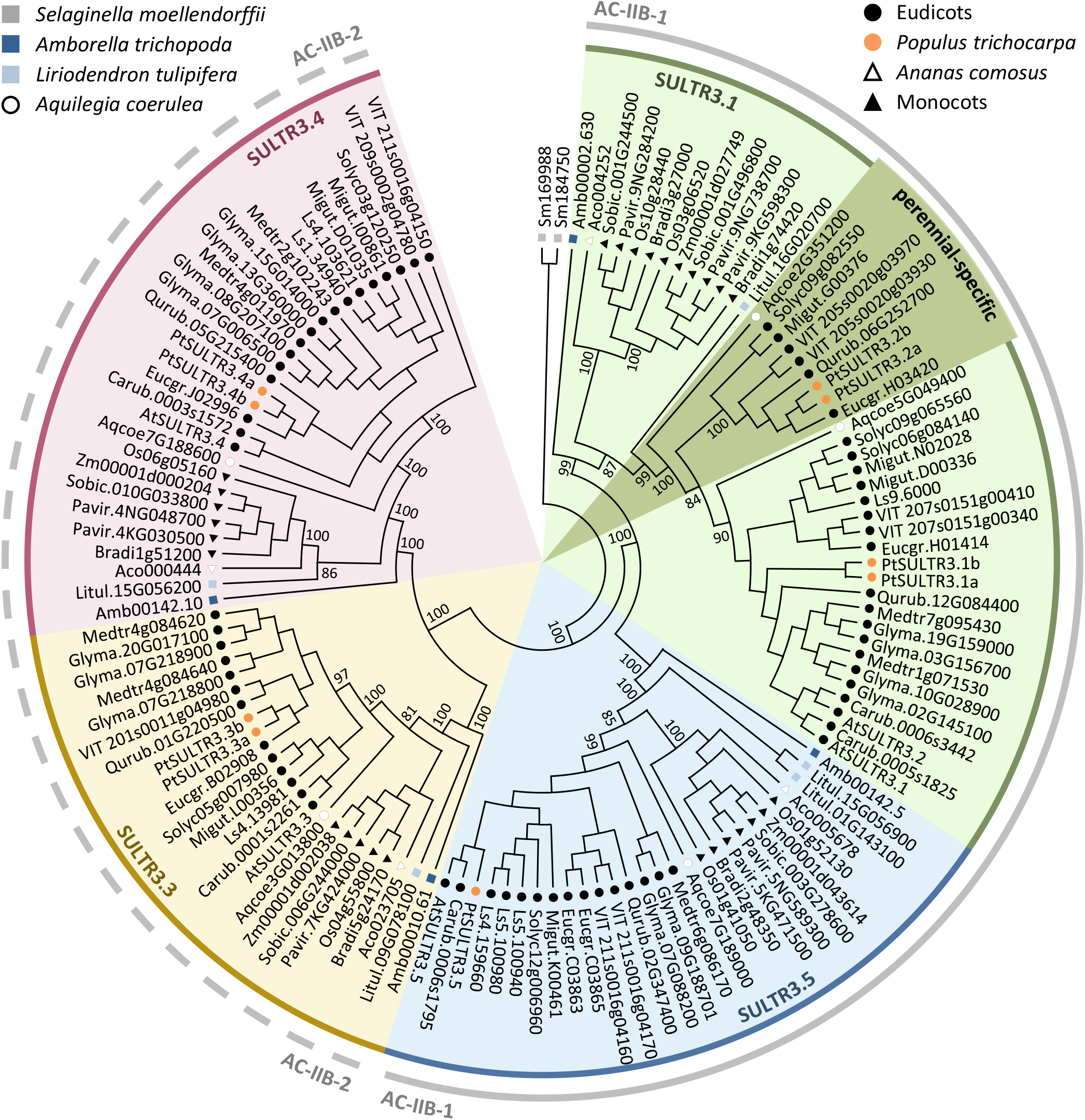
Maximum likelihood phylogenetic inference of AC-IIB members from 20 angiosperm genomes. The 125 angiosperm SULTR3 members form two branches each with two nodes, rooted on the two *Selaginella moellendorffi* sequences. The four color-coded nodes represent SULTR3.1 and SULTR3.5 of AC-IIB-1, and SULTR3.3 and SULTR3.4 of AC-IIB-2. The perennial-specific subgroup within the SULTR3.1 subfamily is highlighted in darker shade. Bootstrap values for major nodes are shown.

### Substrate binding site alignment and protein structural modeling

All poplar and Arabidopsis SULTR protein sequences were aligned as above. The structure-resolved AtSULTR4.1 (AT5G13550) protein sequence was used as the reference to annotate key residues involved in substrate and co-substrate binding and interactions according to Wang et al. (2021). Protein sequences for PtaSULTR3.1a, PtaSULTR3.2a, PtaSULTR3.3a, PtaSULTR3.4a, and PtaSULTR3.5 from *Populus tremula* × *alba* INRA 717-1B4 were used for AlphaFold3 (Abramson et al. 2024) modeling to investigate substrate-binding pocket architecture and potential determinants of substrate specificity. AlphaFold3 models were generated using default parameters (seed 1), without template constraints. Each model was visually inspected for fold quality and overall membrane topology. Structural comparisons and visualizations were performed using PyMOL v3.0.0 (The PyMOL Molecular Graphics System, Schrödinger, LLC).

### Gene expression analysis

RNA-Seq datasets from published *Populus tremula* × *alba* studies (Swamy et al. 2015; Xue et al. 2016; Tsai et al. 2020; Zhou et al. 2023) were obtained from the NCBI Short Read Archive (SRA) using the ‘nfcore/fetchngs’ pipeline (v1.11.0) (Ewels et al. 2020). Quality control and read trimming were performed using Cutadapt v4.4 (Martin 2011) with parameters -q 30, -m 30, and --trim-n. Reads were mapped to the *P. tremula* subgenome (*Populus tremula* × *alba* HAP1 v5.1) of the hybrid (Zhou et al. 2023) available on Phytozome v13. For expression quantification, kallisto v0.48.0 (Bray et al. 2016) was used, with a *k*-mer size of 21 for indexing. Single-end reads were quantified with an estimated average fragment length of 200 and a standard deviation of 20. Paired-end reads were quantified without the stranded option. *SULTR* gene expression values were extracted and statistical significance of treatment responses was performed using PRISM v10.1.0 (GraphPad Software, Boston, MA). Heatmap and bar graphs were generated using Excel.

Gene coexpression analysis was performed using 174 RNA-seq datasets from *Populus tremula* × *alba* stem vascular tissues, excluding the four studies cited above (Swamy et al. 2015; Xue et al. 2016; Tsai et al. 2020; Zhou et al. 2023). The full list of datasets is provided in Supplemental Table S2. Data were similarly processed to obtain expression values. Genes with low expression (TPM < 5 in all samples) were filtered out and only those with a coefficient of variance (CV) > 50% were retained. Pairwise Pearson correlation coefficients (PCC) were calculated between each of the remaining 19,787 genes and either *PtaSULTR3.2a* or *PtaSULTR3.4a* across the 174 samples (Supplemental Table S3) using a python script available at https://github.com/TsailabBioinformatics/sultr-coexpression-utils. Genes with PCC ≥ 0.7 relative to *PtaSULTR3.2a* or *PtaSULTR3.4a* were extracted for Gene Ontology (GO) enrichment using ShinyGO v0.77 (Ge et al. 2020) implemented at http://aspendb.uga.edu/ShinyGO/. Further examination of *SULTR3.2a* and *SULTR3.4a* expression in response to transgenic perturbations of the lignin biosynthetic pathway in *P. trichocarpa* was based on the study by Wang et al. (2018). The list of datasets is provided in Supplemental Table S4.

## RESULTS

### Differential expansion of SULTR subfamilies

A total of 262 SULTR protein sequences from 20 angiosperms and two early-divergent land plant lineages were retrieved from Phytozome v13 (Goodstein et al. 2012) (https://phytozome-next.jgi.doe.gov/) (Table 1). These SULTR proteins are typically annotated based on their Arabidopsis orthologs into four groups, SULTR1 to SULTR4 (Takahashi et al. 2000; Takahashi et al. 2012). The number of sequences per group varies due to differential gene expansion among the species studied. The SULTR4 subfamily consists of one or two sequences in each sampled taxon, while the other three groups show varied expansions. SULTR3 has the highest number of sequences, followed by SULTR1, whereas SULTR2 exhibits a modest expansion, observed only in some eudicot species (Table 1).

**Table 1.**
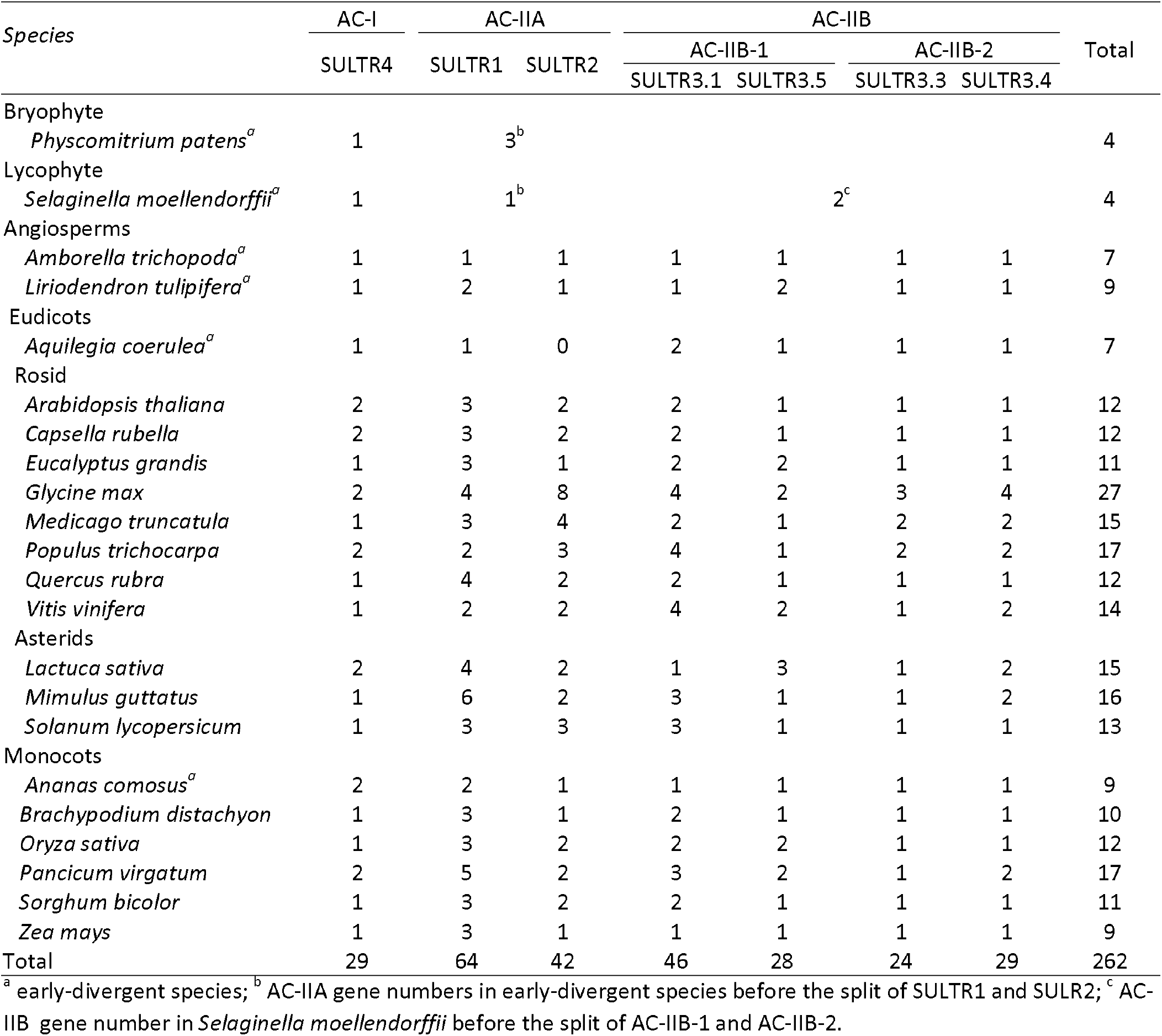
SULTR family members in 22 plant species.

| Species | AC-I | AC-IIA |  | AC-IIB |  |  |  | Total |
| --- | --- | --- | --- | --- | --- | --- | --- | --- |
|  | SULTR4 | SULTR1 | SULTR2 | AC-IIB-1 |  | AC-IIB-2 |  |  |
|  |  |  |  | SULTR3.1 | SULTR3.5 | SULTR3.3 | SULTR3.4 |  |
| Bryophyte |  |  |  |  |  |  |  |  |
| *Physcomitrium patens*<sup>a</sup> | 1 |  | 3<sup>b</sup> |  |  |  |  | 4 |
| Lycophyte |  |  |  |  |  |  |  |  |
| *Selaginella moellendorffii*<sup>a</sup> | 1 |  | 1<sup>b</sup> |  |  | 2<sup>c</sup> |  | 4 |
| Angiosperms |  |  |  |  |  |  |  |  |
| *Amborella trichopoda*<sup>a</sup> | 1 | 1 | 1 | 1 | 1 | 1 | 1 | 7 |
| *Liriodendron tulipifera*<sup>a</sup> | 1 | 2 | 1 | 1 | 2 | 1 | 1 | 9 |
| Eudicots |  |  |  |  |  |  |  |  |
| *Aquilegia coerulea*<sup>a</sup> | 1 | 1 | 0 | 2 | 1 | 1 | 1 | 7 |
| Rosid |  |  |  |  |  |  |  |  |
| *Arabidopsis thaliana* | 2 | 3 | 2 | 2 | 1 | 1 | 1 | 12 |
| *Capsella rubella* | 2 | 3 | 2 | 2 | 1 | 1 | 1 | 12 |
| *Eucalyptus grandis* | 1 | 3 | 1 | 2 | 2 | 1 | 1 | 11 |
| *Glycine max* | 2 | 4 | 8 | 4 | 2 | 3 | 4 | 27 |
| *Medicago truncatula* | 1 | 3 | 4 | 2 | 1 | 2 | 2 | 15 |
| *Populus trichocarpa* | 2 | 2 | 3 | 4 | 1 | 2 | 2 | 17 |
| *Quercus rubra* | 1 | 4 | 2 | 2 | 1 | 1 | 1 | 12 |
| *Vitis vinifera* | 1 | 2 | 2 | 4 | 2 | 1 | 2 | 14 |
| Asterids |  |  |  |  |  |  |  |  |
| *Lactuca sativa* | 2 | 4 | 2 | 1 | 3 | 1 | 2 | 15 |
| *Mimulus guttatus* | 1 | 6 | 2 | 3 | 1 | 1 | 2 | 16 |
| *Solanum lycopersicum* | 1 | 3 | 3 | 3 | 1 | 1 | 1 | 13 |
| Monocots |  |  |  |  |  |  |  |  |
| *Ananas comosus*<sup>a</sup> | 2 | 2 | 1 | 1 | 1 | 1 | 1 | 9 |
| *Brachypodium distachyon* | 1 | 3 | 1 | 2 | 1 | 1 | 1 | 10 |
| *Oryza sativa* | 1 | 3 | 2 | 2 | 2 | 1 | 1 | 12 |
| *Panicum virgatum* | 2 | 5 | 2 | 3 | 2 | 1 | 2 | 17 |
| *Sorghum bicolor* | 1 | 3 | 2 | 2 | 1 | 1 | 1 | 11 |
| *Zea mays* | 1 | 3 | 1 | 1 | 1 | 1 | 1 | 9 |
| Total | 29 | 64 | 42 | 46 | 28 | 24 | 29 | 262 |

<sup>a</sup> early-divergent species; <sup>b</sup> AC-IIA gene numbers in early-divergent species before the split of SULTR1 and SULTR2; <sup>c</sup> AC-IIB gene number in *Selaginella moellendorffii* before the split of AC-IIB-1 and AC-IIB-2.

We constructed a SULTR phylogenetic tree using maximum likelihood inference with IQ-TREE (Nguyen et al. 2015) (Figure 1). The SULTR members formed distinct clusters in a topology consistent with previous reports (Takahashi et al. 2012; Ding et al. 2016; Zhao et al. 2022; Chen et al. 2024). This topology suggests the existence of two ancient clades, based on the placement of SULTRs from the bryophyte moss (*Physcomitrella patens*), as also suggested by Takahashi et al. (2012). The smaller Ancient Clade I (AC-I) comprises the tonoplast SULTR4, whereas the larger Ancient Clade II (AC-II) encompasses the other three SULTR groups (Figure 1). Within AC-II, the ancient vascular plant spikemoss (*Selaginella moellendorffii*) contains two distinct groups that are basal to two subclades: one represented by SULTR1-SULTR2 (referred to as AC-IIA) and the other by SULTR3 (AC-IIB). This suggests that the divergence of the two subclades coincides with the evolution of vascular plants (Figure 1).

### SULTR1 and SULTR2 originate from ancient local duplication

AC-IIA consists of two strongly supported nodes represented by the SULTR1 and SULTR2 subfamilies. Both subfamilies are evolutionarily conserved in angiosperms, including the basal lineages Amborella (*Amborella trichopoda*) and tulip tree (*Liriodendron tulipifera*). Interestingly, in both genomes, *SULTR1* and *SULTR2* are found in close proximity with a divergent (head-to-head) orientation. This head-to-head pattern has been maintained in nearly all sampled angiosperm taxa, despite lineage-specific variations, including additional whole-genome or tandem duplications (see below, Supplemental Table S1). This suggests that *SULTR1* and *SULTR2* originated from a local duplication event before the advent of angiosperms. The data further indicates that the proximally located and oppositely oriented *SULTR1* and *SULTR2* copies are likely the founding members of each subfamily.

Monocot SULTR1s formed two clusters, each represented by all sampled taxa, including the basal lineage *Ananas comosus* (pineapple) (Figure 1). This suggests their divergence early during monocot evolution. Eudicot SULTR1s also formed two distinct clusters. The smaller cluster has limited species representation, and there is only a single SULTR1 in the basal eudicot blue columbine (*Aquilegia coerulea*)(Figure 1). This suggests that the expansion of SULTR1 in eudicots occurred later and only in certain taxa. In the smaller eudicot SULTR1 cluster, four of the eight species—eucalyptus (*Eucalyptus grandis*), lettuce (*Lactuca sativa*), monkey flower (*Mimulus guttatus*), and oak (*Quercus rubra*)—harbor tandem duplicates (Supplemental Table S1). This suggests a tendency for lineage-specific local amplification of this SULTR1 subgroup.

The SULTR2 subfamily is smaller than SULTR1, with 1-2 copies of the two basal angiosperm lineages and monocots (Figure 1, Table 1). However, the SULTR2 subfamily size is more variable among eudicots, ranging from a complete absence in blue columbine to four to eight copies in the legumes (Table 1). In the latter case, we detected whole genome, tandem, and other duplication events both before and after the split of barrelclover and soybean, leading to a disproportionately expanded SULTR2 subfamily in these lineages (Supplemental Table S1).

### SULTR3 encompasses distinct subfamilies predating angiosperm evolution

AC-IIB encompasses the entire SULTR3 group, with spikemoss members at the basal position (Figure 1). This supports the vascular-specific origin of SULTR3s as previously suggested (Takahashi et al. 2012). Like AC-IIA, AC-IIB is divided into two well-supported nodes, each containing two distinct branches. The first node, hereafter AC-IIB-1, includes the SULTR3.1/3.2 and SULTR3.5 branches, while the second node, AC-IIB-2, comprises the SULTR3.3 and SULTR3.4 branches (Figure 1). Each of these four branches contains one or two members from Amborella and tulip tree, which are sister to all flowering plants (Table 1). This indicates that, like SULTR1 and SULTR2, the four SULTR3 branches evolved before the emergence of angiosperms.

To better understand the evolutionary history of AC-IIB, we constructed a phylogenetic tree using only SULTR3 sequences (Figure 2). The overall topology mirrors that of the AC-IIB group in Figure 1, with angiosperm members clustering into four branches predating the emergence of Amborella and tulip tree. Three of the four branches correspond to the SULTR3.3, SULTR3.4, and SULTR3.5 subfamilies, each with distinct monocot and dicot groups consistent with their early divergence during angiosperm evolution (Figure 2). Most species are represented by a single copy, while others harbor two to four paralogs (Table 1).

The SULTR3.1 branch (also called SULTR3.1/3.2 branch in previous papers, see below) has substantially expanded in both monocots and eudicots after their split. Within the monocots, we identified two strongly supported subgroups for grass SULTR3.1s, rooted with the lone member of pineapple (Figure 2), indicating their origin from a grass-specific duplication event (Figure 2). We also observed two distinct subgroups for eudicots, including blue columbine, indicating their divergence predates the common ancestor of eudicots (Figure 2). The larger subgroup includes all 12 sampled eudicot taxa, many of which have additional paralogs due to lineage-specific duplications. Examples include Brassicaceae (Arabidopsis AtSULTR3.1 and AtSULTR3.2), Salicoid (poplar PtSULTR3.1a and PtSULTR3.1b), and legume-specific duplications (Figure 2). In contrast, the smaller eudicot subgroup is only retained in blue columbine and six other sampled taxa in both rosids and asterids, all of which are perennial species (Figure 2). Distinct orthologs were also identified in other perennial eudicot genomes available on Phytozome v13, such as birch (*Betula platyphylla*), papaya (*Carica papaya*), chestnut (*Castanea dentata*), orange (*Citrus sinensis*), cassava (*Manihot esculenta*), peach (*Prunus persica*), and castor bean (*Ricinus communis*) of rosid, and coffee (*Coffea arabica*) and hydrangea (*Hydrangea quercifolia*) of asterids. These findings suggest that this SULTR3.1 subgroup may play an important role in the functionality of perennial species.

The duplication history of the SULTR3.1 branch has led to inconsistent gene and subfamily nomenclatures in the literature. Specifically, Arabidopsis AtSULTR3.1 and AtSULTR3.2 are paralogs within the same SULTR3.1 (large) subgroup, unlike AtSULTR3.3, AtSULTR3.4, and AtSULTR3.5 which represent phylogenetically distinct subfamilies. In poplar, PtSULTR3.1 and PtSULTR3.2 are descendants of the ancestral eudicot duplication in the larger and smaller SULTR3.1 subgroups, respectively, with their Salicoid paralogs designated with “a” and “b” suffixes (*e.g*., PtSULTR3.1a and PtSULTR3.1b) in the literature (Dürr et al. 2010; Malcheska et al. 2017). In these examples, AtSULTR3.1/3.2 are orthologous to PtSULTR3.1a/b, while the perennial-specific PtSULTR3.2a/b do not have direct orthologs in Arabidopsis (Figure 2). We preserve the use of SULTR3.2 as a gene name in keeping with convention and refer to this subfamily as SULTR3.1 for now. Phylogenomics-informed gene (re)naming is clearly needed but is beyond the scope of this study (see Tweedie et al. 2025). Our analysis strongly supports four distinct SULTR3 (AC-IIB) subfamilies SULTR3.1, SULTR3.5 (node AC-IIB-1), SULTR3.3, and SULTR3.4 (node AC-IIB-2), alongside the two, well studied AC-IIA subfamilies SULTR1 and SULTR2. Together with the more divergent AC-I SULTR4 group, angiosperm taxa contain seven evolutionarily conserved SULTR subfamilies (Figure 1), rather than the historically proposed four (Takahashi et al. 2000).

### Exon-intron structure varies with the evolution of SULTR subfamilies

Analysis of gene structure revealed variations that generally align with the SULTR subfamily evolution inferred from our phylogenetic trees. *SULTR4* genes of the small AC-I clade have 17 exons with evolutionarily conserved junctions among diverse lineages (Figure 3). However, multiple exon-fusion events have occurred in the expanded AC-II subfamilies. Specifically, three fusions between exons 8 and 9, exons 10 to 12, and exons 13 and 14 are shared across all AC-II subfamilies and species, including basal spikemoss members (Figure 3). Among them, *Sm184750* (basal to AC-IIB) is likely the founding member with just these three exon fusions. This gene structure is retained in *SULTR1* (AC-IIA), *SULTR3.3* and *SULTR3.4* (AC-IIB-2) members.

**Figure 3.**
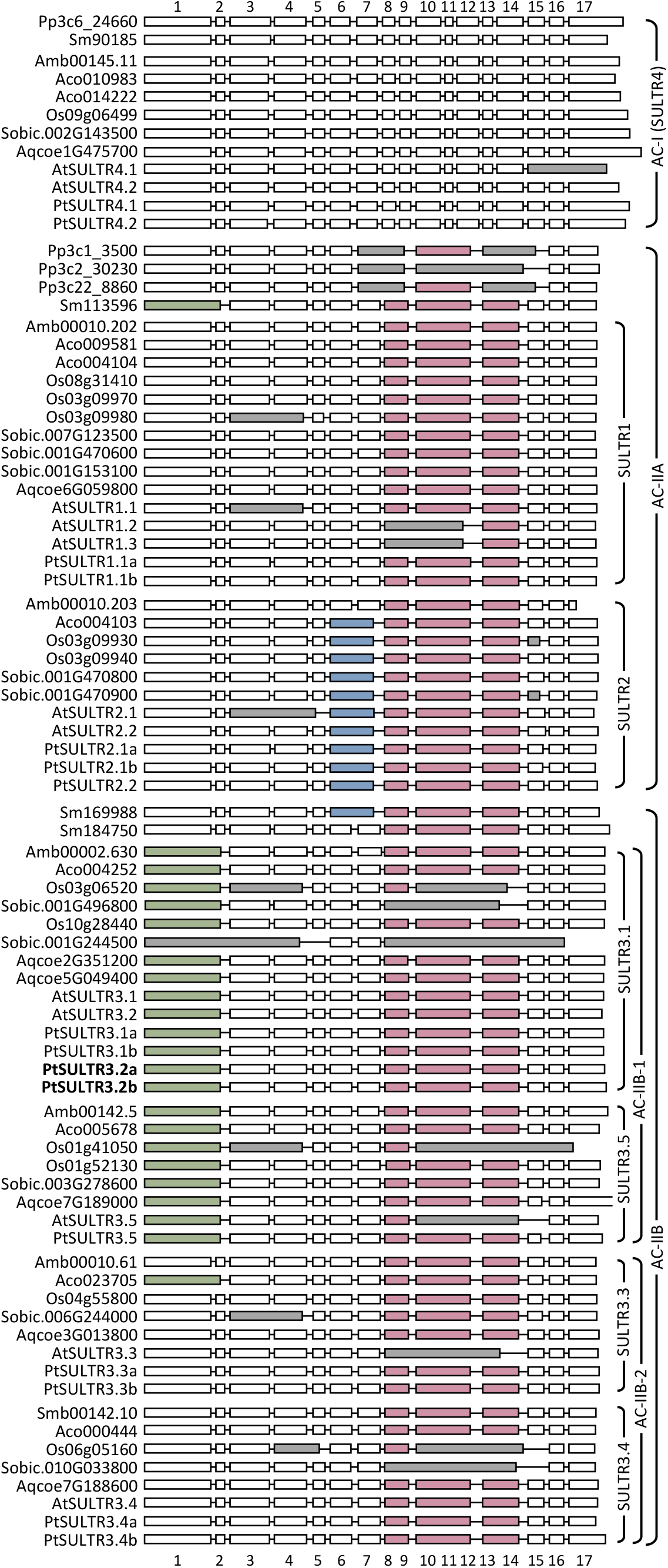
Exon-intron structures of *SULTR* genes of nine representative species. Genes are arranged by phylogenetic classification, with basal lineages shown on top of each group. Conserved exon fusion events are colored and gray denotes lineage-specific fusion events. Introns are not drawn to scale and first exons are trimmed for alignments. Boldface indicates perennial-specific SULTR3.1 subfamily members.

*SULTR2* members (AC-IIA) have an additional fusion between exons 6 and 7. This fusion is absent in the basal Amborella *SULTR2* which retains the same gene structure as the presumed AC-II founding member *Sm184750* and the sister *SULTR1* branch (Figure 3). Interestingly, this fusion is present in the more distantly related spikemoss *Sm169988* basal to AC-IIB (Figure 3). Similarly, the *SULTR3.1* and *SULTR3.5* subfamilies share a conserved fusion of exons 1 and 2 across basal, monocot, and eudicot lineages of the AC-IIB-1 subclade, as well as the less related *Sm113596* basal to AC-IIA (Figure 3). The independent occurrence of the same exon fusion pattern in unrelated lineages may suggest convergent evolution. Alternatively, the minor inconsistencies noted above may reflect limitations in current phylogenetic reconstructions in correctly positioning distant orthologs from early-divergent lineages, such as moss and spikemoss. This limitation is evident for the SULTR4 clade (Figure 1) as previously reported (Takahashi et al. 2012).

Finally, gene structure analysis also revealed multiple unusual or lineage-specific fusion events. Examples include grass-specific fusions, including secondary fusions between above-mentioned exon fusion events, in *SULTR3.1* and *SULTR3.4* members (Figure 3, Phytozome v13). Furthermore, half of the Arabidopsis *SULTR* family members have unique exon fusion events conserved in Brassicaceae, indicative of their unusual evolution. This suggests that using Arabidopsis genes as sole references for comparative studies may introduce confounding factors for evolutionary inference.

### Protein modeling reveals binding pocket substitutions in expanded SULTR3 subfamilies

The recently resolved protein structure of Arabidopsis AtSULTR4.1 (Wang et al. 2021) was used as a template to identify key substrate binding residues in other Arabidopsis and poplar SULTR proteins. The key residue Glu347 (AtSULTR4.1 numbering), essential for co-substrate proton sensing and transport, is conserved across all SULTRs (Figure 4A). Key residues lining the sulfate binding pocket, including Gln112, Tyr116, Ser392, and Arg(Lys)393, are also conserved (Figure 4A). Mutations to Ala at any of these positions resulted in a near-total loss of sulfate transport activity (Wang et al. 2021). However, two residues within the predicted binding pocket showed subfamily-specific substitutions. Ala153 is replaced by Ser in SULTR3.3 and SULTR3.4 (node AC-IIB-2), while Ser390 is replaced by Pro in SULTR3.1 and SULTR3.5 (node AC-IIB-1) or by Ala in SULTR3.3 (Figure 4A). Ser390 is thought to interact with Arg393, which likely provides positive electrostatic potential to stabilize substrate binding (Wang et al. 2021). Whether these substitutions contribute to the reported difficulty in demonstrating *in vitro* sulfate transport activity for SULTR3s (Takahashi et al. 2000; Kataoka et al. 2004a) or to the neofunctionalization of SULTR3.4 as SPDT for phosphate transport (Yamaji et al. 2017; Ding et al. 2020) is a possibility that merits consideration.

**Figure 4.**
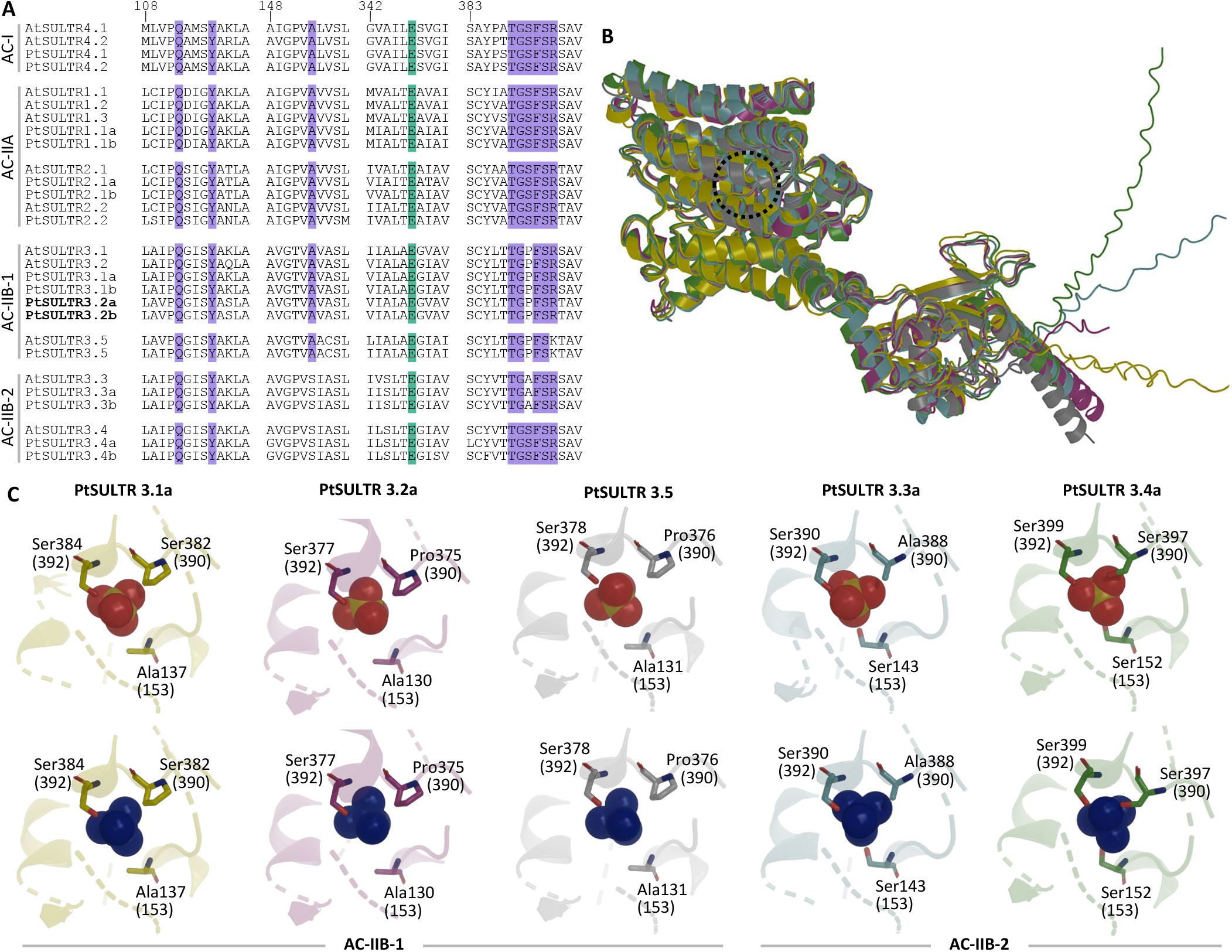
Sequence and structural comparisons of SULTR members. ***A***, Sequence alignments of *Arabidopsis thaliana* and *Populus trichocarpa* SULTRs near the predicted substrate-binding pocket. Amino acid residue numbering follows the structure-resolved AtSULTR4.1 (AT5G13550), shown at the top. Other sequences are arranged by phylogenetic grouping. Boldface indicates perennial-specific SULTR3.1 subfamily members. Green- and purple-shaded positions denote residues critical to proton and sulfate binding or interaction, respectively. ***B***, Structure alignment of AlphaFold3 models of PtaSULTR3.1a (yellow), PtaSULTR3.2a (purple), PtaSULTR3.3a (blue), PtaSULTR3.4a (green), and PtaSULTR3.5 (grey) showing overall fold conservation. The dotted circle denotes the substrate-binding site. ***C***, Close-up views of the substrate-binding pockets docked with sulfate (SO4^2-^, red spheres, top row) or phosphate (PO_4_^3-^, blue spheres, bottom row). Key residues are highlighted, with AtSULTR4 numbering shown in parentheses.

To explore this further, we performed AlphaFold3 (Abramson et al. 2024) structural modeling for five representative poplar AC-IIB members: PtaSULTR3.1a, PtaSULTR3.2a, PtaSULTR3.5 (AC-IIB-1), PtaSULTR3.3a, and PtaSULTR3.4a (AC-IIB-2). Structural alignments revealed conserved overall folding across AC-IIB members (Figure 4B). Most models accommodated sulfate (SO4^2□^) and phosphate (PO4^3□^) ions in similar positions (Figure 4C). However, SULTR3.4a displayed a distinct binding mode, with PO4^3□^positioned differently from SO4^2□^, suggesting potential structural selectivity. The Ala→Ser153 substitution in AC-IIB-2 introduces a polar hydroxyl group capable of hydrogen bonding. In SULTR3.4a, this residue, along with Ser390 and Ser392, forms a triad of serines that interact with the docked PO4^3□^ (Figure 4C), which may help stabilize its higher negative charge (™3). Although SULTR3.3 also contains Ser153, its Ser→Ala390 substitution likely disrupts favorable PO4^3□^ interactions. The other AC-IIB-1 SULTR3 members possess only one Ser residue proximal to the substrate, suggesting suboptimal coordination at the binding site (Figure 4C).

### Expression divergence and tissue partitioning in expanded SULTR3 subfamilies

We analyzed the expression of poplar *SULTR* genes using published RNA-Seq datasets, including the JGI tissue atlas from *P. tremula* × *P. alba* INRA 717-1B4 (Zhou et al. 2023). The AC-I *PtaSULTR4* transcripts were detected in all tissues, with *PtaSULTR4.1* being higher than *PtaSULTR4.2* in mature leaves and young elongating roots (Figure 5). Among the AC-IIA members, the two *PtaSULTR1.1* paralogs were expressed in all tissues but showed distinct tissue preferences. *PtaSULTR1.1a* transcript levels were highest in the bark and xylem of the primary stem (internodes 3-6), while *PtaSULTR1.1b* transcripts were low overall but predominated in roots (Figure 5). The *PtaSULTR2.1* paralogs were generally poorly expressed, except in primary xylem where *PtaSULTR2.1b* was strongly expressed. *PtaSULTR2.2*, the ancient inverted tandem duplicate of *PtaSULTR1.1b*, was well expressed throughout the plant with a clear bias for mature leaves (Figure 5).

**Figure 5.**
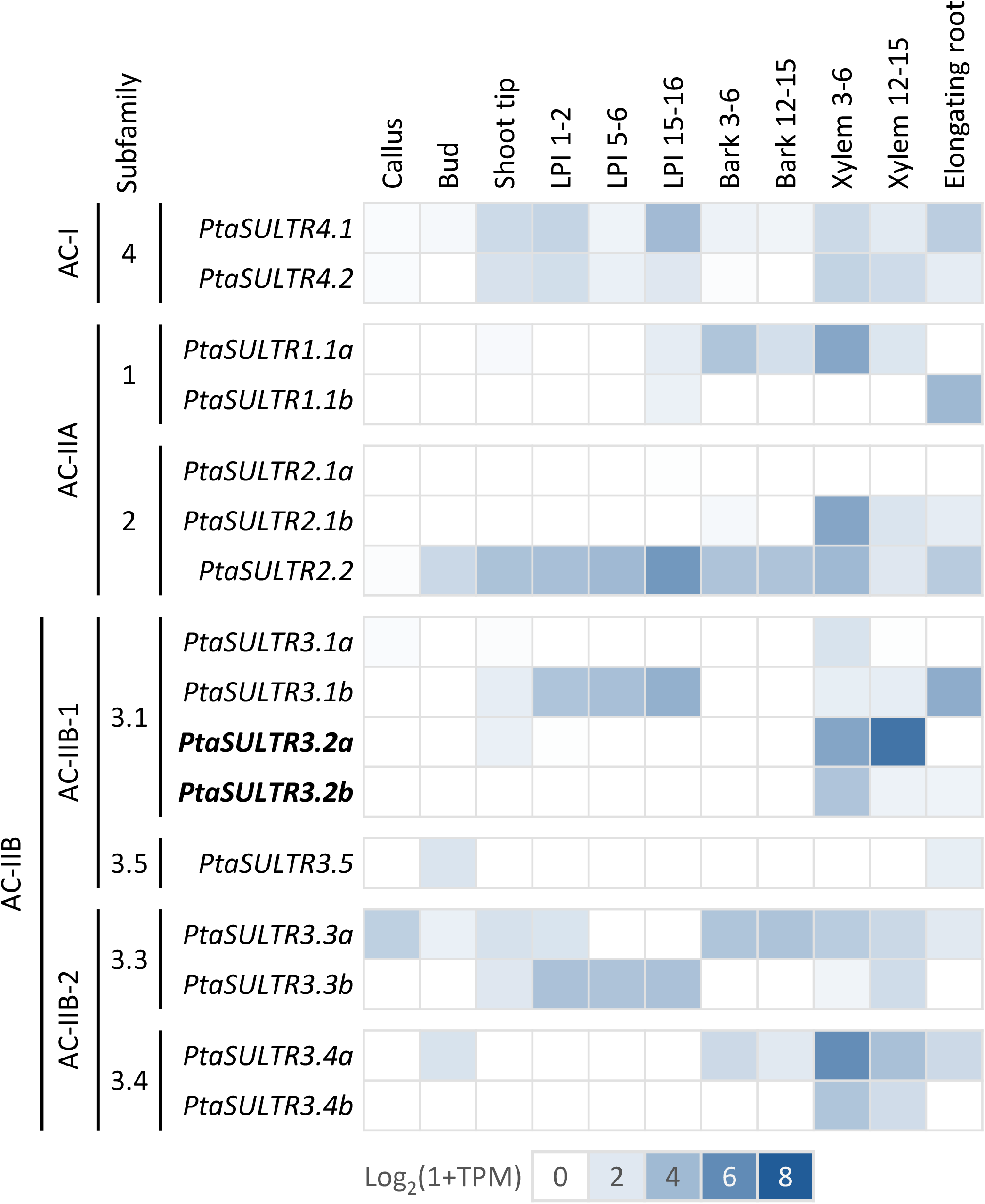
Heatmap illustration of *SULTR* gene expression across *Populus* tissues. Genes are arranged by phylogenetic grouping and genome duplicates are denoted as a and b. Boldface indicates perennial-specific SULTR3.1 subfamily members. Data were from JGI tissue atlas of Zhou et al. 2023.

Members of the expanded AC-IIB subfamilies exhibited substantial expression divergence or partitioning, with many showing restricted tissue distribution. For instance, the singleton *PtaSULTR3.5* was only detected at low levels in buds and roots (Figure 5). No bark expression was detected for the *PtaSULTR3.1* and *PtaSULTR3.5* (AC-IIB-1) subfamilies. *PtaSULTR3.1a* was poorly expressed overall, but the paralogous *PtaSULTR3.1b* was well expressed in both leaves and roots (Figure 5). The *PtaSULTR3.2* pair showed clearly xylem-biased and developmentally conditioned expression in stems. Both genes were expressed in primary xylem, but *PtaSULTR3.2a* expression increased substantially in secondary xylem as *PtaSULTR3.2b* expression sharply decreased (Figure 5). Primary xylem-biased expression was also detected for the *PtaSULTR3.4* paralogs, with *PtaSULTR3.4a* being higher than *PtaSULTR3.4b. PtaSULTR3.3* is the only AC-IIB subfamily that showed broad tissue expression, but this expression was partitioned between the paralogs. *PtaSULTR3.3a* was mainly expressed in stem and root tissues, whereas *PtaSULTR3.3b* was detected in leaves (Figure 5).

### Differential responses of xylem-biased SULTR3.2a and SULTR3.4a to drought and lignification

Some *PtaSULTR* genes have previously been shown to be sensitive to seasonal changes in wood or to drought stress in leaves and roots based on qRT-PCR studies (Dürr et al. 2010; Malcheska et al. 2013; Malcheska et al. 2017). Here, we explored data from stress and transgenic perturbation RNA-Seq experiments (Xue et al. 2016; Tsai et al. 2020; Swamy et al. 2015) to analyze *SULTR* expression responses, focusing particularly on vascular tissues and the expanded *SULTR3* subfamilies (excluding *PtaSULTR3.5* which was below detection). We observed a partitioned drought response both between subfamilies and between paralogs, consistent with the partitioned tissue preference discussed above (Figures 5-6). The dominant *PtaSULTR3.1* subfamily members, *PtaSULTR3.1b* and *PtaSULTR3.2a*, were significantly upregulated by drought in roots and secondary xylem, respectively, while their poorly expressed paralogs were unresponsive (Figure 6). Similar patterns were observed for the *PtaSULTR3.3* and *PtaSULTR3.4* subfamilies, with the dominant *PtaSULTR3.3a* and *PtaSULTR3.4a* significantly upregulated in drought-stressed bark and root, respectively. Although both *PtaSULTR3.1b* and *PtaSULTR3.4a* were drought-responsive in root tissue, the expression of *PtaSULTR3.1b* was an order of magnitude higher than that of *PtaSULTR3.4a*. In summary, the data showed that each of the three vascular tissues responded to drought stress by upregulating a distinct AC-IIB (*SULTR3*) subfamily member: *PtaSULTR3.1b* in roots, *PtaSULTR3.2a* in xylem, and *PtaSULTR3.3a* in bark.

**Figure 6.**
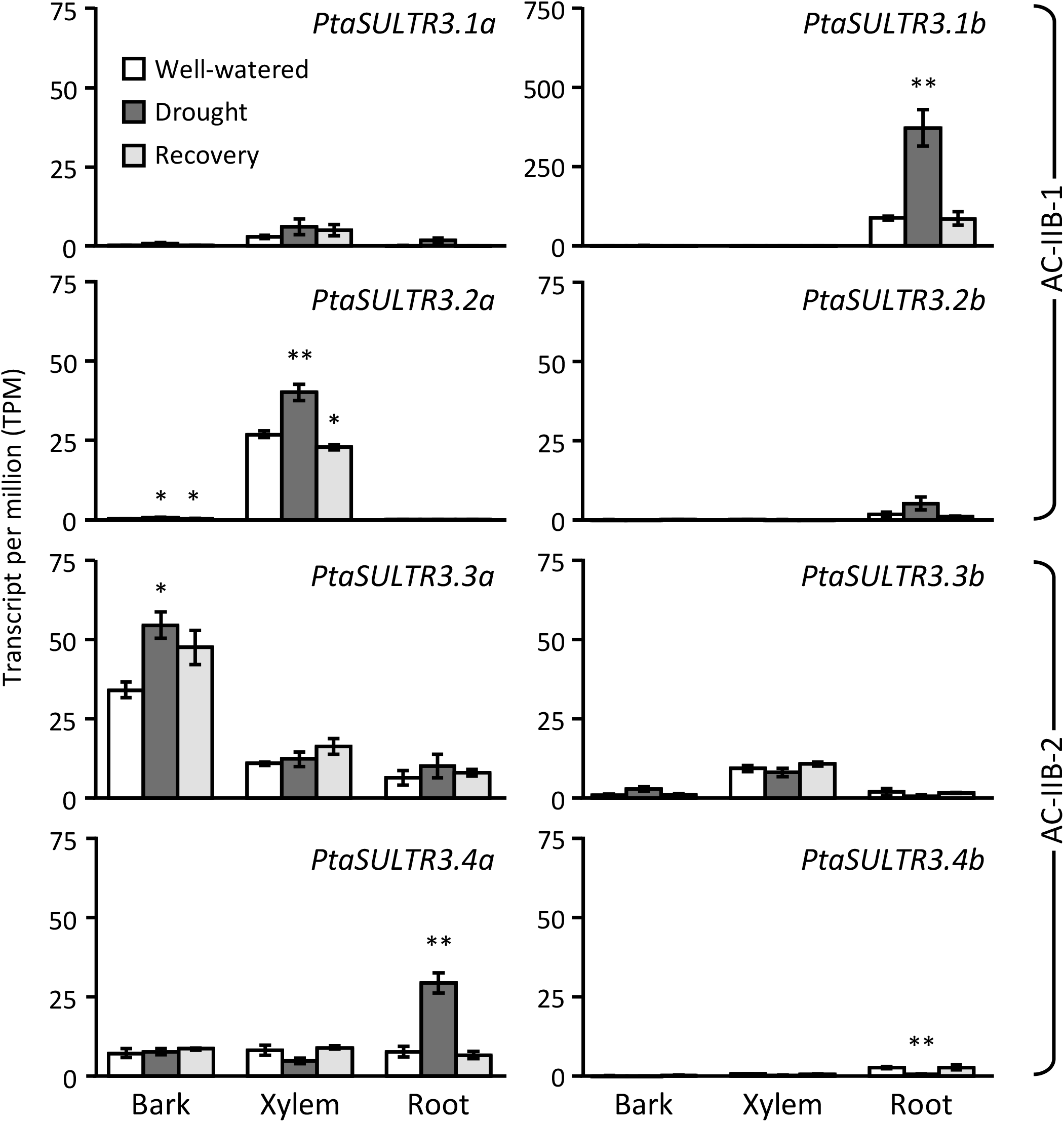
Drought stress response of *Populus SULTR3* members. The four *SULTR3* subfamilies showed partitioned tissue response to drought stress, with divergent expression strengths between genome duplicates. Bars are means ± SE of n = 3 biological replicates. Significance testing was conducted using the two-sample *t*-test against well-watered samples (**, *P* ≤ 0.01; * *P* ≤ 0.05). Data were from Xue et al. 2016.

We next examined *SULTR3* expression in the secondary xylem (internodes 60-80) of erect or inclined poplar trees during normal wood (NW) or tension wood (TW) development (Swamy et al. 2015). The secondary xylem-predominant *PtaSULTR3.2a* was down-regulated by more than 70% in TW (Figure 7A). TW in angiosperms is characterized by cellulose-enriched gelatinous fibers with overall reduced lignin accrual (Groover 2016). To further investigate the potential connection between *SULTR3* expression and cell wall composition, we utilized RNA-Seq data from young developing xylem (15 to 30 cm from the shoot tip) of *4CL1* (*4-coumarate:CoA ligase*)-knockout poplar with reduced lignin accumulation (Zhou et al. 2015; Tsai et al. 2020). We observed a 40-50% downregulation of both *PtaSULTR3.2a* and *PtaSULTR3.4a* in the low-lignin *4cl1* mutants (Figure 7B). Interestingly, *PtaSULTR3.4a* transcript abundance was higher than that of *PtaSULTR3.2a*, presumably reflecting the younger stems used in this study compared to the TW experiment (Swamy et al. 2015; Tsai et al. 2020). Together, these results suggest that *PtaSULTR3.2a* expression correlated positively with lignification during stem development, while *PtaSULTR3.4a* response was conditional and limited to primary xylem, where it was most highly expressed (Figures 5 and 7).

**Figure 7.**
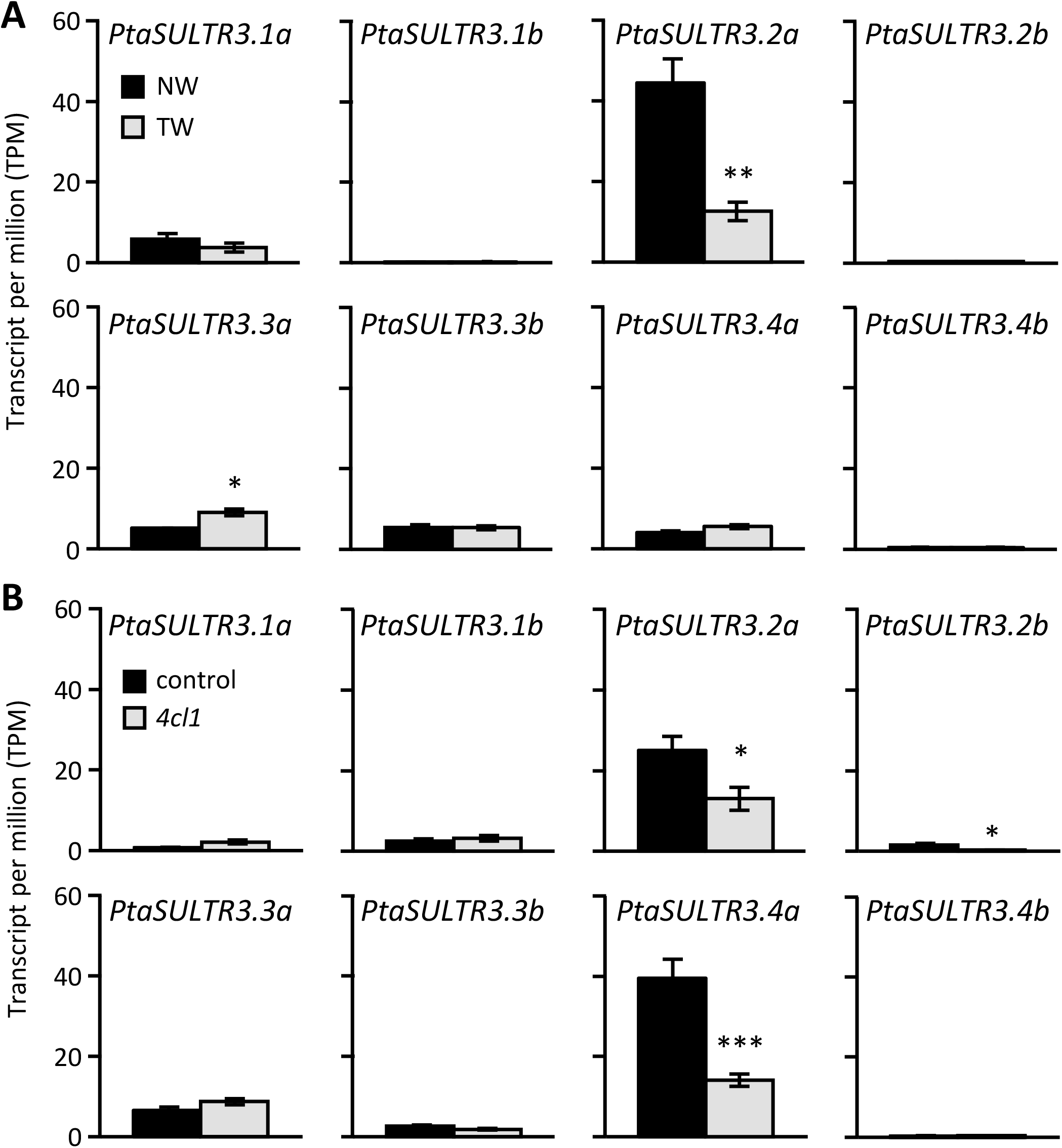
Xylem expression responses of *Populus SULTR3* members to genetic or environmental perturbations. *A*, Transcript abundance in normal wood (NW) versus tension wood (TW) from *P. tremula* × *alba* stem internodes 60-80 (Swamy et al. 2015). *B*, Transcript abundance in *P. tremula* × *alba* control versus lignin-reduced *4cl1* xylem of young stem internodes 15-30 cm from the shoot tip (Tsai et al. 2020). Bars are means ± SE of n = 3 biological replicates. Statistical significance was assessed using two-sample *t*-test (***, *P* ≤ 0.001; **, *P* ≤ 0.01; * *P* ≤ 0.05).

### Gene coexpression of PtaSULTR3.4a with phosphorus starvation response and PtaSULTR3.2a with lignification

To further explore the functional association of the xylem-biased *PtaSULTR3.2a* and *PtaSULTR3.4a*, we performed gene-centered coexpression analysis using 174 published RNA-seq datasets from xylem, bark, and stem tissues (Supplemental Table S2). We identified 419 genes coexpressed with *PtaSULTR3.4a* (Pearson correlation coefficient or PCC ≥ 0.7), which showed significant enrichment in Gene Ontology (GO) terms associated with “cellular response to phosphate starvation”, “sulfolipid biosynthetic process”, “response to nitrate”, “sulfate transport”, and “phosphate ion transport” (Figure 8A). Indeed, *PtaSULTR3.4a* was strongly coexpressed with *PtaSULTR3.4b* (PCC = 0.91), *PtaSULTR2.1b* (0.82), and *PtaSULTR2.2* (0.72), as well as genes encoding multiple phosphate transporters (Supplemental Table S3), consistent with previous reports linking the SULTR3.4 subfamily to phosphorus allocation (Yamaji et al. 2017; Ding et al. 2020). *PtaSULTR3.4a* was also highly coexpressed with genes known to be sensitive to phosphorus starvation, including orthologs of SPX domain-containing proteins and purple acid phosphatases (Supplemental Table S3) (Duan et al. 2008; Wang et al. 2011; Gan et al. 2015; Chen et al. 2022). Our analysis suggests a close coordination of *PtaSULTR3.4a* with a network of transporters and phosphorus responses genes.

**Figure 8.**
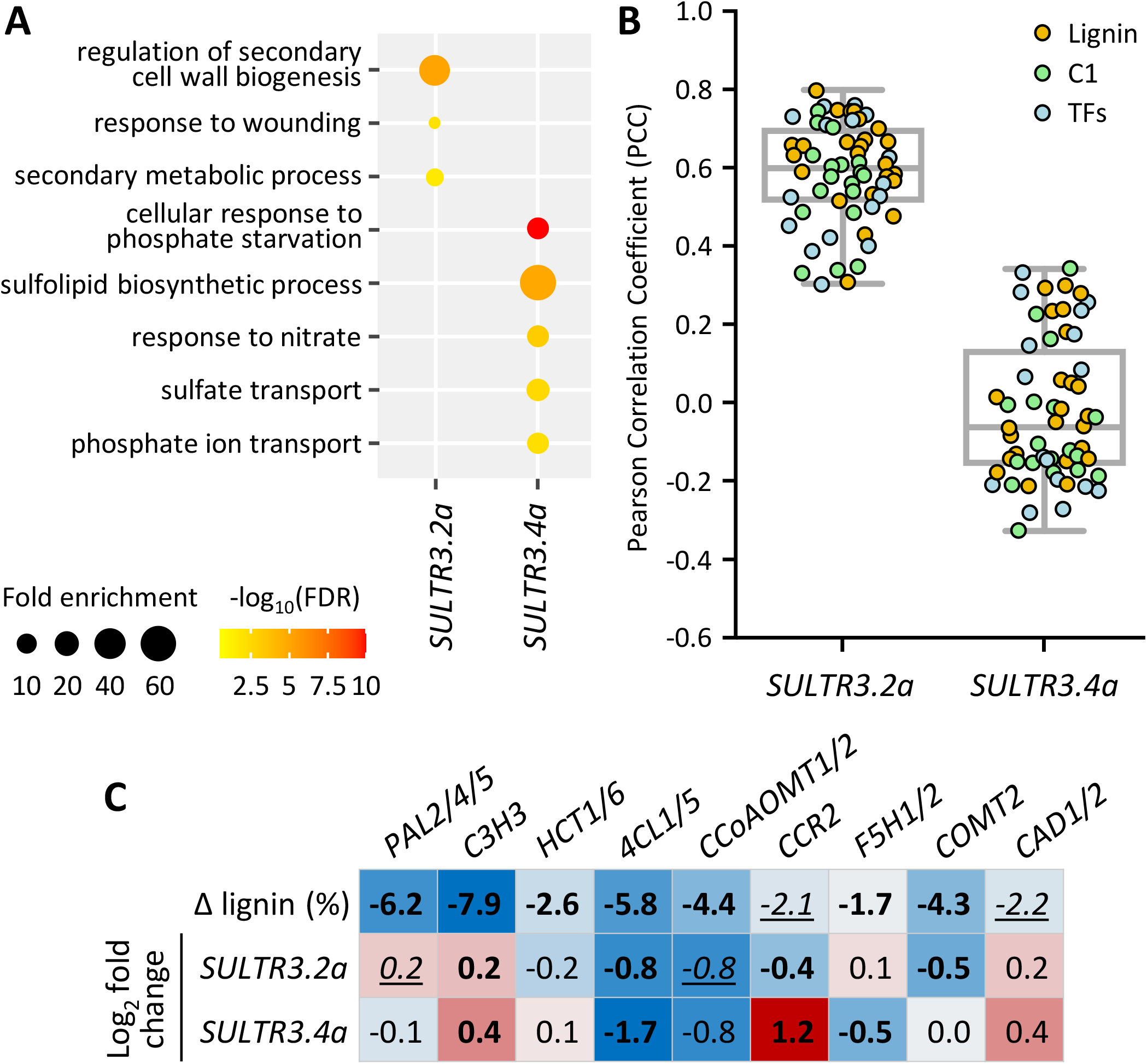
Gene-centered coexpression analysis of xylem-biased *PtaSULT3.2a* and *PtaSULTR3.4a*. ***A***, GO enrichment of 87 and 419 genes co-expressed (Pearson correlation coefficient [PCC] ≥ 0.7) with *PtaSULT3.2a* and *PtaSULTR3.4a*, respectively, across 174 published *P. tremula* × *alba* RNA-seq datasets from stem vascular tissues. ***B***, Boxplot distribution of PCC values against 58 genes annotated and color-coded for their involvement in lignin biosynthesis, C1 metabolism, and upstream transcription factors (TFs). The gene list is provided in Supplemental Table S3. ***C***, Heatmap depictions of lignin content differentials and expression fold changes of *SULTR3.2a* and *SULTR3.4a* in transgenic *P. trichocarpa* with downregulated lignin biosynthetic genes (Wang et al. 2018). Statistical significance was determined using two-sample *t*-test (boldfaced, *P* ≤11.01; italic-underlined, *P* ≤11.05).

In contrast, a smaller group of 87 genes was coexpressed with *PtaSULTR3.2a*, with significant GO enrichment in “regulation of secondary cell wall biogenesis”, “response to wounding”, and “secondary metabolic process” (Figure 8A). Specifically, *PtaSULTR3.2* was coexpressed with genes encoding multiple lignin biosynthetic enzymes (phenylalanine ammonia lyase [PAL], cinnamate 4-hydroxylase, 4CLs, hydroxycinnamoyl-CoA shikimate/quinate hydroxycinnamoyl transferase, ferulic acid 5-hydroxylase), C1 metabolism enzymes (serine hydroxymethyltransferase [SHMT] and *S*-adenosylmethionine [SAM] synthases), and transcriptional factors regulating secondary wall biogenesis, such as SND (secondary wall-associated NAC domain) and MYB proteins (Zhong et al. 2011) (Supplemental Table S3). We then plotted PCC values for 58 genes involved in lignin biosynthesis, C1 metabolism, and their upstream transcriptional regulators (Supplemental Table S3). We found a median PCC of 0.60 with *PtaSULTR3.2a* but no correlation was observed with *PtaSULTR3.4a* (Figure 8B). Notably, *SULTR3.2a* was positively coexpressed with C1 metabolism genes encoding Met synthases, SAM synthases, homocysteine *S*-methyltransferases, and *S*-adenosylhomocysteine hydrolase of the activated methyl cycle, as well as SHMT and Gly decarboxylase complex enzymes involved in Ser-Gly interconversion (Hanson and Roje 2001)(Figure 8B, Supplemental Table S3). For *PAL* and *4CL* gene families involved in both lignin and flavonoid biosynthesis, *PtaSULTR3.2a* coexpression was observed specifically for lignin-associated *PtaPAL2* (Kao et al. 2002), *Pta4CL1* and *Pta4CL5* isoforms (Harding et al. 2002; Tsai et al. 2020) (Supplemental Table S3).

To further substantiate the link between *SULTR3.2a* and lignification, we mined additional RNA-Seq datasets from transgenic *Populus trichocarpa* with reduced lignin by RNA interference (RNAi) of pathway genes (Wang et al. 2018)(Supplemental Table S4). We confirmed significant down-regulation of *SULTR3.2a* in lignin-reduced transgenic plants, but only for perturbations at the 4CL (consistent with Figure 7B), caffeoyl-CoA *O*-methyltransferase (CCoAOMT), cinnamoyl-CoA reductase (CCR), or caffeic acid *O*-methyltransferase (COMT) steps (Figure 8C). These enzymatic steps involve either CoA-thioester as a co-substrate (4CL) or co-product (CCR) or SAM as a co-factor (CCoAOMT and COMT). Both CoA and SAM are derived from sulfur-containing amino acids (Hanson and Roje 2001; Kupke et al. 2003). In contrast, *SULTR3.4a* was down-regulated in *4CL*-silenced *P. trichocarpa* as in *P. tremula* × *alba* (Figures 7B and 8C), but exhibited variable responses to other lignin gene perturbations, indicative of an indirect connection. Together, independent transgenic experiments in two poplar species and large-scale coexpression analysis all support a previously undescribed link between *SULTR3.2a* and lignification.

## DISCUSSION

The evolutionary expansion and functional diversification of SULTR transporters from bacteria to higher plants has been extensively discussed (Takahashi et al. 2012). However, important questions remain, especially regarding the large and vascular plant-specific SULTR3 group, whose members exhibit little to no sulfate transport activity on their own (Takahashi et al. 2000; Kataoka et al. 2004a). SULTR3 members exhibit variability in their membrane localization and have been reported to facilitate chloroplast sulfate assimilation, enhance sulfate transport by other SULTR partners, and contribute to phosphorus distribution via the vasculature (Kataoka et al. 2004a; Cao et al. 2013; Zhao et al. 2016; Yamaji et al. 2017; Chen et al. 2019; Ding et al. 2020; Gu et al. 2022). By including multiple early-divergent species at key evolutionary junctures for phylogenetic inference, we updated the SULTR family classification into seven distinct subfamilies dating back to basal angiosperms, with four belonging to the enigmatic SULTR3 group. Consistent with experimental evidence from the Arabidopsis, rice, and barley orthologs (Yamaji et al. 2017; Ding et al. 2020; Gu et al. 2022), we provide protein modeling and gene coexpression data supporting the potential neofunctionalization of SULTR3.4 in phosphate homeostasis. We further identified a perennial-specific subgroup within the SULTR3.1 subfamily, with the poplar member *PtaSULTR3.2a* showing strong coexpression with lignin biosynthetic genes and responsiveness to lignin pathway perturbations.

### Early evolution of seven divergent SULTR subfamilies in angiosperms

Our phylogenetic analysis supports several features of the plant SULTR family evolution previously suggested by Takahashi et al. (2012), based on the placement of moss and spikemoss members. These include the split of AC-I (SULTR4) and AC-II (SULTR1-SULTR3) clades early during land plant evolution, and the subsequent split of AC-IIA (SULTR1-SULTR2) and AC-IIB (SULTR3) subclades, with AC-IIB arising specifically in vascular plants (Figure 1). While AC-IIA is further split into the SULTR1 and SULTR2 subfamilies, the much large AC-IIB was previously considered as one (SULTR3) subfamily with four subgroups (Takahashi et al. 2012). However, our phylogenetic reconstruction with basal angiosperm lineages uncovered four strongly supported SULTR3 subfamilies that predate the split of Amborella and tulip tree (Figure 1). Examination of the recently sequenced camphor tree (*Cinnamomum kanehirae* v3) (Chaw et al. 2019) and water lily (*Nymphaea colorata* v1.2)(Zhang et al. 2020) genomes available on Phytozome v13 confirmed the presence of four AC-IIB subfamilies in other early-divergent angiosperm taxa, suggesting their origin in the last common ancestor of all flowering plants.

Gene structure analysis provided further support for distinct evolution of the AC-IIB subfamilies. It is important to note that gene structure changes, such as intron losses, can impact expression and contribute to functional diversification as effectively, if not more so, than mutations within exons (Morello and Breviario 2008). This was evidenced by preferential yet partitioned expression, both among *SULTR3* subfamilies and between gene duplicates, in vascular tissues of poplar (Figure 5). Drought stress stimulated expression of *PtaSULTR3.1b, PtaSULTR3.2a*, and *PtaSULR3.3a* in a root-, xylem-, and bark-biased manner, respectively (Figure 6). These examples serve to illustrate the diverse roles of the four SULTR3 subfamilies beyond the historically established vacuolar (SULTR4), high-affinity (SULTR1), and low-affinity (SULTR2) sulfate transport activities. The four SULTR3 subfamilies likely differentiated as mechanisms for internal distribution and utilization of sulfate evolved in angiosperms (Takahashi 2019).

### Protein modeling insights into substrate binding pocket sequences

Despite differences in tissue expression and transport kinetics, the proton and sulfate binding pocket residues predicted to be essential for activity (Wang et al. 2021) are conserved in AC-I SULTR4 and AC-IIA SULTR1/SULTR2 subfamilies (Figure 4). However, the expanded AC-IIB (SULTR3) members exhibit subfamily-specific substitutions within the binding pocket, coinciding with reported catalytic divergence. Among AC-IIB members, only SULTR3.1 and SULTR3.5 have been reported to exhibit sulfate uptake activity in yeast, although these findings lack kinetic characterization (Krusell et al. 2005; Xun et al. 2021) and have not been consistently observed across tested species (Takahashi et al. 2000). For instance, Arabidopsis AtSULTR3.5 was found to be nonfunctional by itself but can enhance the activity of AtSULTR2.1 significantly (Kataoka et al. 2004a). Reverse genetics analysis revealed that mutations to AtSULTR3.1 were more detrimental to sulfate uptake by isolated chloroplasts than mutations to any other AC-IIB members (Cao et al. 2013; Chen et al. 2019). Mutations in another SULTR3.1 subfamily member, AtSULTR3.2, has been linked to altered sulfate translocation in developing seeds of Arabidopsis (Zuber et al. 2010). SULTR3.1 and SULTR3.5 differ from the canonical AC-I and AC-IIA subfamilies at position 390 (Pro vs. Ser). Ser390 is thought to interact with Arg393 to stabilize the sulfate ion electrostatically within the binding pocket (Wang et al. 2021). The Ser→Pro390 substitution may impair sulfate ion coordination, potentially contributing to the poor sulfate transport of SULTR3.1 and SULTR3.5 (Takahashi et al. 2000; Kataoka et al. 2004a).

AC-IIB-2 subfamilies SULTR3.3 and SULTR3.4 differ from the other SULTRs at position 153, where Ala is replaced by Ser in the substrate-binding pocket (Figure 4). Furthermore, Ser390 is conserved in SULTR3.4 but replaced by Ala in SULTR3.3. In SULT3.4, the three serine residues—Ser153, Ser390, and Ser392—are predicted to enable PO4^3□^ interaction based on protein-substrate modeling (Figure 4). Multiple studies have shown that SULTR3.4 subfamily members transport phosphate rather than sulfate (Yamaji et al. 2017; Ding et al. 2020; Gu et al. 2022). Interestingly, phosphate ion coordination in the binding pocket is disrupted in the SULTR3.3 model due to the Ser→Ala390 substitution (Figure 4), consistent with the lack of *in vitro* phosphate or sulfate transport activity reported for rice OsSULTR3.3 (Zhao et al. 2016). However, mutations in *OsSULTR3.3* reduced phytate and total phosphorus concentrations in rice grains (Zhao et al. 2016), suggesting a noncanonical function. Together, these findings highlight unique structural adaptations in SULTR3.4/SPDT that support PO4^3□^ interaction, providing a mechanistic basis for its proposed neofunctionalization in phosphate allocation within the vasculature (Yamaji et al. 2017; Ding et al. 2020). The data also hint at the importance of Ser153 and Ser390 for substrate selectivity. Whether the Pro and Ala substitutions at position 390 contribute to the functional diversification of other SULTR3 subfamilies, either as nonfunctional transporter-like facilitators or through noncanonical roles, remains to be investigated experimentally.

### *PtaSULTR3*.4 coexpression with phosphorus transport and starvation response genes

Poplar *PtaSULTR3.4a* and its less expressed genome duplicate *PtaSULTR3.4b* were preferentially expressed in primary xylem. Of the seven Salicoid genome duplicate pairs in the SULTR family, *PtaSULTR3.4s* are the only pair that retains expression conservation in stem vascular tissues. Consistent with a role of the SULTR3.4/SPDT subfamily in phosphorus distribution and metabolism (Yamaji et al. 2017; Ding et al. 2020), *PtaSULTR3.4a*-coexpressed genes included several phosphate transporters and phosphate starvation response markers (Gan et al. 2015; Chen et al. 2022). This group also included genes encoding nitrate-responsive regulators, transporters for nitrate, amino acids, auxin, and various metal ions, as well as plasma membrane H^+^-ATPases necessary to energize transporters for transmembrane nutrient uptake (Palmgren 2001). The coordinated acquisition of nitrogen and phosphorus, and the crosstalk between nitrate- and phosphorus-responsive signaling pathways have been extensively reported, including in poplar (Gan et al. 2015; Hu and Chu 2020; Ueda et al. 2020). This coordination also extends to other mineral nutrients (Raddatz et al. 2020; Wei et al. 2023). Similarly, sulfur assimilation is sensitive to availability of not only sulfate but also nitrate (Kopriva et al. 2004). Our gene coexpression analysis implicated SULTR3.4 as an important player in these coordinated responses.

### Secondary xylem-biased PtaSULTR3.2a expression and lignification

Extensive prior work has illustrated temporal associations between *SULTR* gene expression and sulfate circulation in poplar stems throughout the year (Rennenberg et al. 2007; Dürr et al. 2010; Malcheska et al. 2013). The sulfate assimilation pathway interfaces with C1 metabolism for the production of SAM, which is a methyl donor for the biosynthesis of lignin and other cell wall polysaccharides (Rajinikanth et al. 2007; Srivastava et al. 2015; Zhang et al. 2019). However, there is little information linking SULTRs with wood formation. A connection was recently suggested via a xylem-expressed microRNA, *miR395c*, which regulated sulfate assimilation genes and low-affinity SULTR2.1b to affect ABA synthesis and secondary cell wall biosynthesis in poplar (Liu et al. 2022).

In poplar, the AC-IIB-1 member *SULTR3.2a* is the dominant *SULTR* transcript in secondary xylem undergoing secondary cell wall thickening. A functional association between PtaSULTR3.2a and lignification is supported by its reduced expression in low-lignin xylem, either via genetic perturbation in *4cl1* mutants or in TW of trees subjected to stem bending (Figure 7). Significantly lower *SULTR3.2a* expression was also detected in transgenic RNAi *P. trichocarpa* (Wang et al. 2018) targeting 4CL, CCR, CCoAOMT and COMT—enzymatic steps involving sulfur-containing CoA or SAM (Figure 8C). In addition, xylem *PtaSULTR3.2a* expression increased during drought stress, a condition generally associated with enhanced lignification (Moura et al. 2010). We found strong coexpression of *PtaSULTR3.2a* with genes involved in lignin biosynthesis, C1 metabolism, and secondary cell wall biogenesis (Rajinikanth et al. 2007; Zhong et al. 2011; Zhang et al. 2019; Donev et al. 2023), accounting for 17% of the 87 *PtaSULTR3.2a*-coexpressed genes. While strictly correlative, the conclusion that PtaSULTR3.2a is more closely associated with lignification than other PtaSULTRs is supported by data from a wide range of experimental approaches (Supplemental Table S2), including transgenic perturbation of lignin genes in two distinct species (*P. trichocarpa* of section Tacamahaca and *P. tremula* × *alba* of section *Populus*). Given that PtaSULT3.2a belongs to a small subgroup of the SULTR3.1 subfamily retained only in perennial dicots, its preferential expression in secondary xylem and strong coexpression with lignification genes are consistent with perennial-specific functional diversification.

In summary, our analysis suggests that the enigmatic SULTR3 contains four distinct subfamilies with an ancient origin predating angiosperm evolution. Variations in exon-intron structure, substrate binding pocket residues, and tissue expression preferences likely contribute to the functional diversification of the four SULTR3 subfamilies. The previously reported neofunctionalization of the SULTR3.4 subfamily associated with phosphorus distribution (Yamaji et al. 2017; Ding et al. 2020; Gu et al. 2022) gained credence in poplar based on structural modeling and coexpression analysis. In addition, we provided evidence supporting the functional diversification of PtaSULTR3.2a associated with lignification in poplar and likely other perennial species. These functional predictions await future experimental verification.Finally, this work highlights the need for phylogeny-informed gene and gene family nomenclatures as discussed elsewhere in this special issue.

## ACKNOWLEDGEMENTS

The authors thank the advice of Ethan Baldwin and Jim Leebens-Mack on phylogenetic analysis and Ran Zhou on bioinformatics analysis.

## AUTHORS’ CONTRIBUTIONS

C.-J.T. conceived the idea, S.M.S. performed all analyses with assistance from C.H. for RNA-seq and gene coexpression, L.N. performed protein modeling analysis, S.M.S. drafted the manuscript, S.A.H. and C.-J.T. revised the manuscript.

## SUPPLEMENTARY DATA

Table S1. List of SULTR sequences used in the phylogenetic analysis

Table S2. List of RNA-Seq datasets from *Populus tremula* × *alba* stem tissues used in coexpression analysis

Table S3. Genes and their Pearson correlation coefficients with *PtaSULTR3.2a* and *PtaSULTR3.4a* Table S4. List of RNA-Seq datasets from transgenic *Populus trichocarpa*

## DATA AVAILABILITY STATEMENT

Gene models and genome versions for protein sequences used for phylogenetic analysis are provided in Supplementary Table S1. Accession numbers for RNA-Seq datasets used in gene expression analysis are SRR6255832 to SRR6255842 for tissue expression, RR1292611, SRR1292615, and SRR1295980-SRR1295983 for stem bending response, SRP041959 for drought response, and PRJNA589632 for lignin perturbation response in *P. tremula* × *alba*. Accession numbers for the 174 RNA-Seq datasets used for gene coexpression analysis are provided in Supplemental Table S2. The accession number for additional RNA-Seq data from lignin-reduced transgenic *P. trichocarpa* is PRJNA314500 and individual sample accession numbers are provided in Supplemental Table S4.

